# p66Shc Mediates SUMO2-induced Endothelial Dysfunction

**DOI:** 10.1101/2024.01.24.577109

**Authors:** Jitendra Kumar, Shravan K. Uppulapu, Sujata Kumari, Kanika Sharma, William Paradee, Ravi Prakash Yadav, Vikas Kumar, Santosh Kumar

**Affiliations:** Department of Internal Medicine, Division of Cardiovascular Medicine, University of Iowa, Iowa City, IA; Department of Genetics, Cell Biology and Anatomy, University of Nebraska Medical Center, Omaha, NE; Genome Editing Core Facility, University of Iowa, Iowa City, IA; Regional Institute of Education-NCERT, Mysore, Karnataka, India; Department of Biochemistry and Molecular Biotechnology, Mass Spectrometry Facility, University of Massachusetts Chan Medical School, Worcester, MA

**Author notes:** **Corresponding author** Santosh Kumar, PhD, University of Iowa, Department of Internal Medicine, Division of Cardiovascular Medicine, François M. Abboud Cardiovascular Research Center, 25 South Grand Ave., ML B191, Iowa City, IA-52242, **Email:**. **Author Contributions:** JK, SKU, S Kumari, KS, CA, S Kumar performed studies, analyzed data, and prepared figures; WP and RPY contributed to reagent generation and validation; VK and S Kumar analyzed data, prepared figures and wrote manuscript.

**Keywords:** Sumoylation, endothelial function, p66Shc, SUMO

## Abstract

**Background:** Sumoylation is a post-translational modification that can regulate different physiological functions. Increased sumoylation, specifically conjugation of SUMO2/3 (small ubiquitin-like modifier 2/3), is detrimental to vascular health. However, the molecular mechanism mediating this effect is poorly understood.

**Methods:** We used cell-based assays and mass spectrometry to show that p66Shc is a direct target of SUMO2 and SUMO2 regulates p66Shc function via lysine-81 modification. To determine the effects of SUMO2-p66ShcK81 on vascular function, we generated p66ShcK81R knockin mice and crossbred to LDLr^-/-^ mice to induce hyperlipidemia. Next, to determine p66ShcK81-SUMO2ylation-induced changes in endothelial cell signaling, we performed mass spectrometry followed by Ingenuity Pathway Analysis.

**Results:** Our data reveal that p66Shc mediates the effects of SUMO2 on endothelial cells. Mass spectrometry identified that SUMO2 modified lysine-81 in the unique collagen homology-2 domain of p66Shc. SUMO2ylation of p66Shc increased phosphorylation at serine-36, causing it to translocate to the mitochondria, a step critical for oxidative function of p66Shc. Notably, sumoylation-deficient p66Shc (p66ShcK81R) was resistant to SUMO2-induced p66ShcS36 phosphorylation and mitochondrial translocation. P66ShcK81R knockin mice were resistant to endothelial dysfunction induced by SUMO2ylation and hyperlipidemia. Ingenuity Pathway Analysis revealed multiple signaling pathways regulated by p66ShcK81-SUMO2ylation in endothelial cells, highlighting JAK-STAT as a major pathway affected by SUMO2-p66ShcK81.

**Conclusions:** Collectively, our work reveals SUMO2-p66Shc signaling as a fundamental regulator of vascular endothelial function. We discovered that p66ShcK81 is an upstream modification regulating p66Shc signaling and mediates hyperlipidemia-induced endothelial dysfunction and oxidative stress.

## Introduction

Endothelial cells play an important role in regulating vascular health. One of the primary functions of endothelial cells is to maintain vascular tone and regulate blood flow in response to physiological changes. In the context of disease, endothelial cells either fail to produce enough nitric oxide (NO) or the available NO is quenched by reactive oxygen species (ROS). Consequently, this leads to endothelial dysfunction (ED), characterized by the reduced availability of NO and impaired vasorelaxation.^1^ ED is associated with various conditions such as diabetes, hyperlipidemia, hyperglycemia, obesity, and aging.^2^ There is a strong correlation between ED and atherosclerotic changes in vessel walls.^3^ Therefore, ED is one of the prime targets to prevent pathologies including myocardial infarction and stroke.

Sumoylation is a dynamic post-translational modification involved in many pathologies including vascular disease.^4^ Increased sumoylation promotes ED and subsequent development of atherosclerosis.^5, 6^ Sumoylation occurs by conjugation of **s**mall **u**biquitin-like **mo**difier**s** (SUMOs) to the lysine moiety of the target protein, which may change its fate and/or function. SUMO conjugation is predicted to occur at a specific motif, ΨKXE, where Ψ represents an aliphatic-branched amino acid, X a random amino acid, and E glutamic acid, although other non-canonical sumoylation sites have also been reported.^4, 7, 8^ There are four SUMO isoforms (SUMO1, SUMO2, SUMO3 and SUMO4) and among them SUMO2 is the most abundantly expressed. Unlike SUMO1 and SUMO3, genetic deletion of SUMO2 is embryonically lethal.^9, 10^ While sumoylation is known to promote cardiovascular diseases, the effect of SUMO isoforms on the vasculature is not well studied. Recently, we reported that endothelial-specific overexpression of SUMO2 impairs vascular endothelial function in mice.^11^ However, the molecular mechanisms mediating the endothelial effects of SUMO2 are not known.

p66Shc is a member of the ShcA family of adaptor proteins. Among the ShcA protein family (p66Shc, p52Shc, and p46Shc), p66Shc is the largest protein due to an additional collagen homology domain (CH2) at its N-terminus. Posttranslational modifications in the CH2 domain regulate the oxidative function of p66Shc;^12-15^ whereas phosphorylation in the CH1 domain leads to mitogenic signaling.^16, 17^ p66Shc is abundantly expressed in endothelium and mediates ED due to hyperlipidemia, hyperglycemia, and aging.^18^ Correspondingly, mice with a genetic deletion of p66Shc have a longer life span and are protected against ED due to metabolic disorders.^19, 20^ Here, we examined whether p66Shc mediates the effects of SUMO2 on endothelial cells. Using multiple techniques, we show that SUMO2 directly modifies p66Shc at lysine-81 in its CH2 domain, which is essential for SUMO2-induced ROS production and ED. Thus, our study uncovers a previously unknown posttranslational modification that regulates p66Shc function in the endothelium and defines a novel molecular mechanism by which increased SUMO2ylation induces ROS, leading to vascular ED.

## Results

### p66Shc mediates SUMO2-induced ROS production in endothelial cells

To address whether p66Shc mediates SUMO2-induced ROS production, we overexpressed SUMO2 in human umbilical vein endothelial cells (HUVECs) with and without knockdown of p66Shc and used MitoSOX to detect mitochondrial level of ROS, a prominent function of p66Shc. As expected, overexpression of SUMO2 led to a robust increase in levels of ROS (Fig. 1A and B). This SUMO2-induced increase in ROS was significantly blunted in HUVECs with p66Shc knockdown (Fig. 1A and C). In a complementary approach, we used H_2_DCFDA to determine cellular ROS level and noted a similar increase in ROS level with overexpression of SUMO2 which was significantly reduced by the knockdown of p66Shc (Fig. 1D).This suggests that SUMO2 is targeting p66Shc for ROS production in endothelial cells.

**Figure 1:**
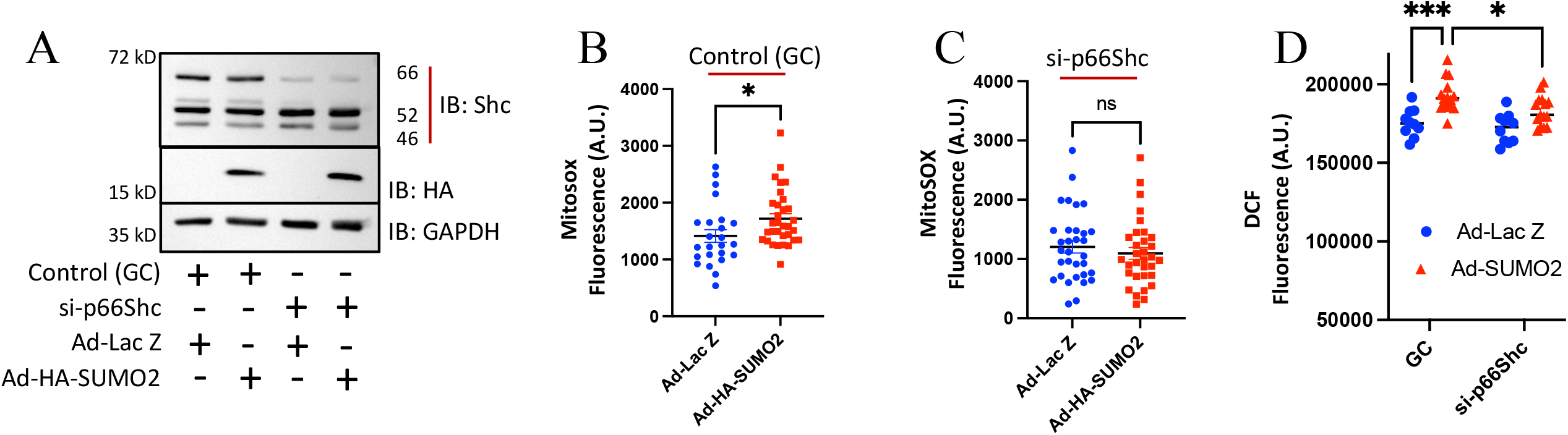
p66Shc mediates SUMO2-induced ROS production and inflammation in endothelial cells. (A) Immunoblot for Shc and SUMO2 expression in HUVECs following p66Shc knockdown (si-p66Shc) and SUMO2 overexpression (Ad-HA-SUMO2). (B and C) Quantification of ROS (MitoSOX fluorescence) in HUVECs from A. *P<0.05; n=24-32; student’s t-test. (D) Quantification of ROS in HUVECs following p66Shc knockdown (si-p66Shc) and SUMO2 overexpression (Ad-HA-SUMO2). ***P<0.001, P<0.05; n=10-15; One-way ANOVA. Data represents mean ±SEM. Immunoblots are representative of at least three independent experiments. GC, scrambled siRNA used as control; HUVECs, human umbilical vein endothelial cells.

### p66Shc is directly modified by SUMO2

We next asked whether p66Shc is a direct target for SUMO2. We overexpressed p66Shc in HEK-293 cells with and without SUMO2. Immunoprecipitation of p66Shc showed many slower migrating protein bands when SUMO2 and p66Shc were co-expressed, suggesting that p66Shc may be SUMO2ylated (Fig. 2A). To confirm that the slower migrating bands were due to sumoylation, we overexpressed SUMO-ligase (Ubc9) to facilitate sumoylation, and a deSUMOylating enzyme (SENP1) which removes SUMO proteins. Overexpression of Ubc9 increased the signal of slower migrating bands whereas SENP1 reduced the signal suggesting the slower migrating bands were due to sumoylation (Fig. 2B). Next, we asked whether the SUMO2-p66Shc interaction exists in endothelial cells by overexpressing SUMO2 and p66Shc in HUVECs. Immunoprecipitation of flag-tagged p66Shc revealed sumoylation (Fig. 2C) which consistently increased in intensity with increasing SUMO2 expression (Fig. S1). To verify that p66Shc undergoes SUMO2ylation in HUVECs we treated cells with a pharmacological inhibitor of sumoylation (anacardic acid). This reduced sumoylation of p66Shc (Fig. 2D &E), demonstrating that indeed p66Shc undergoes SUMO2ylation in endothelial cells. We also performed an *in vitro* sumoylation of p66Shc using recombinant His-tagged p66Shc protein and a commercially available kit (Enzo, Farmingdale, NY), which showed that recombinant p66Shc is modified by SUMO2 (Fig. 2F). Collectively, these findings indicate that p66Shc is a direct target of SUMO2.

**Figure 2:**
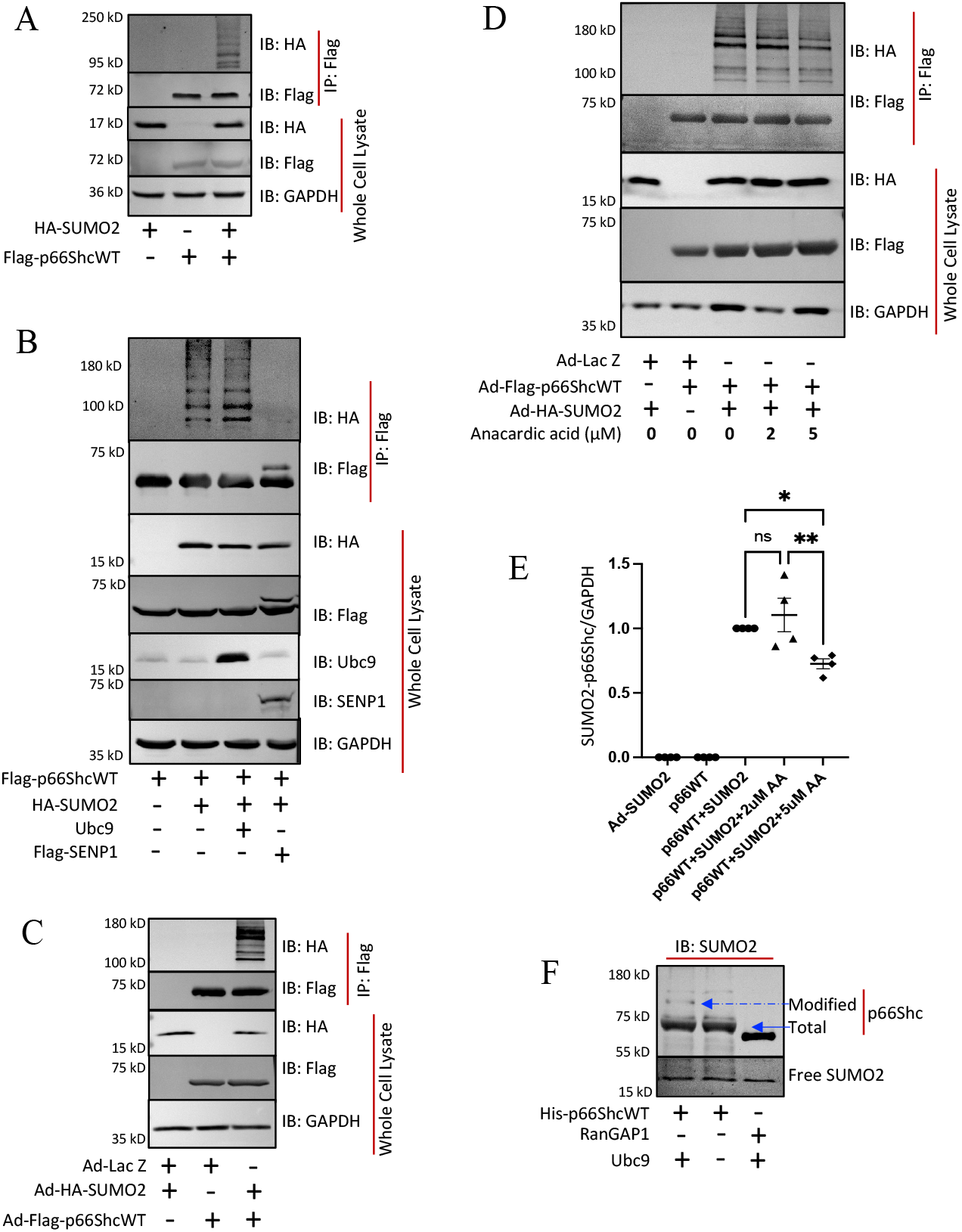
SUMO2 directly modifies p66Shc. (A) Immunoblot for sumoylated p66Shc in HEK-293 cells overexpressing SUMO2 (HA-SUMO2) and p66Shc (Flag-p66Shc). (B) Immunoblot for p66Shc sumoylation in HEK-293 cells overexpressing SUMO2 (HA-SUMO2) and p66Shc (Flag-p66Shc) with or without overexpression of Ubc9 (HA-Ubc9) and SENP1 (Flag-SENP1). (C) Immunoblot for sumoylation of p66Shc in HUVECs overexpressing SUMO2 (Ad-HA-SUMO2) or p66Shc (Ad-Flag-p66Shc). (D) Immunoblot for p66Shc sumoylation in HUVEC cells overexpressing SUMO2 (Ad-HA-SUMO2) or p66Shc (Ad-Flag-p66Shc) in the presence or absence of anacardic acid. Quantification of SUMO2-p66Shc from immunoblot of D. One-way ANOVA, followed by Tukey’s post-hoc analysis, n=4 *P<0.05, **P<0.01. (F) Immunoblot for sumoylated recombinant p66Shc in the presence of absence of Ubc9. Immunoblots are representative of at least three independent experiments. Data represents mean ±SEM. Ad-Lac Z was used as adenoviral control to compensate for the difference in MOI among the groups.

### SUMO2 modifies p66Shc via lysine-81

The CH2 domain of p66Shc is highly conserved and involved in oxidative function of p66Shc. Within this region there are several conserved lysine residues. Among these is lysine 81, which is part of the ΨKXE motif required for sumoylation (Fig. 3A) Therefore, to examine if K81 is sumoylated, we took advantage of SUMO2ylated recombinant p66Shc and resolved it on SDS-PAGE. Based on the approximate location of modified protein bands seen in immunoblotting, we cut out the gel piece and subjected it to mass spectrometry analysis (Fig. S3). This demonstrated that lysine-81 (K81) in p66Shc was SUMO2ylated, identified as lysine conjugated with the QTGG motif present at c-terminal of SUMO2 (Fig. 3B). To confirm that SUMO2 modifies p66Shc at K81, we mutated this residue to Arginine (p66ShcK81R) and overexpressed it in HEK-293 cells with SUMO2 to determine if sumoylation still occurred. Immunoprecipitation showed that p66ShcK81R is resistant to SUMO2ylation, whereas it still occurred with the wild-type (WT) form (Fig. 3C). Similar results were also observed in HUVECs (Fig. 3D). We next performed an *in vitro* SUMO2ylation of recombinant p66ShcWT and p66ShcK81R. As anticipated, immunoblotting showed that, unlike p66ShcWT, p66ShcK81R is resistant to SUMO2ylation (Fig. 3E). Next, to determine the endogenous SUMO2-p66Shc, we generate a custom antibody against the branched peptide, mimicking the region of SUMO2 attachment (SUMO2-K81) to p66Shc using a commercial source (YenZym, CA). Using this SUMO2-p66ShcK81 antibody, we performed immunoprecipitation and immunoblotting in HUVECs treated with o-LDL (200 μg/ml) for 48 hours and noted p66Shc-SUMO2ylation at endogenous level (Fig. 3F). Thus, our findings demonstrate that SUMO2 modifies p66Shc at K81.

**Figure 3:**
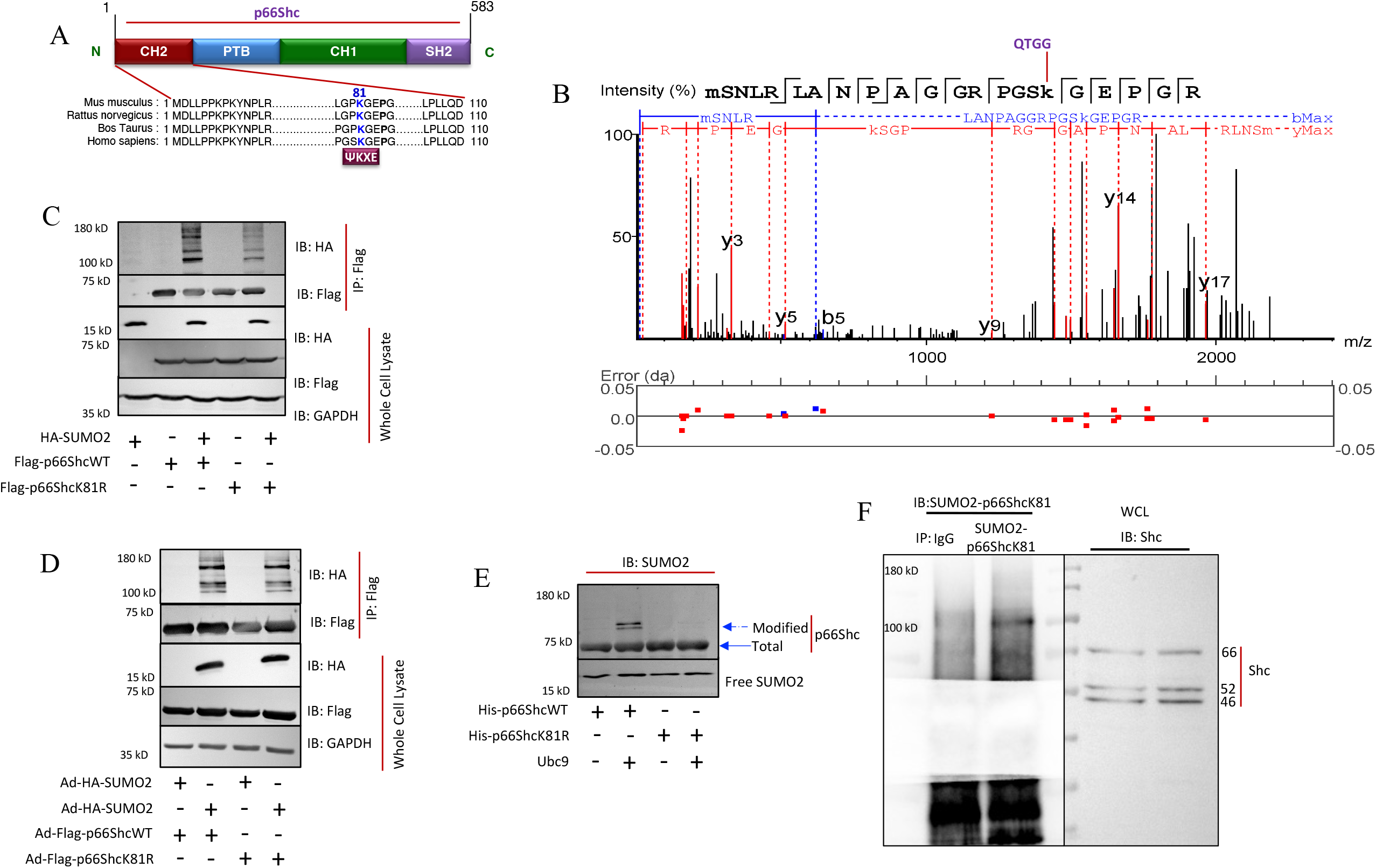
SUMO2 modifies p66Shc at lysine-81. (A) Modular structure of p66Shc showing conserved lysine-81 in the CH2 domain. (B) Tandem mass spectrometry of *in vitro SUMO*2ylated recombinant p66Shc. (C) Immunoblot showing sumoylation of p66Shc in HEK-293 cells overexpressing p66ShcWT (Flag-p66ShcWT) or p66ShcK81R (Flag-p66ShcK81R) with and without SUMO2 (HA-SUMO2). (D) Immunoblot showing sumoylation of p66Shc in HUVECs overexpressing p66ShcWT (Ad-Flag-p66ShcWT) or p66ShcK81R (Ad-p66ShcK81R) with and without SUMO2 (Ad-HA-SUMO2). (E) Immunoblot for sumoylation of recombinant p66ShcWT or p66ShcK81R. (F) Immunoprecipitation of SUMO2-p66Shc using SUMO2-p66ShcK81 antibody or the matching IgG control in HUVECs followed by immunoblotting with SUMO2-p66ShcK81 antibody showing endogenous SUMO2ylation of p66Shc (left-side panel). Immunoblotting with Shc antibody in whole cell lysate (right-side panel). Immunoblots are representative of at least three independent experiments. CH2, collagen homology 2; PTB, phosphotyrosine binding; CH1, Collagen homology 1; SH2, Src homology 2.

### SUMO2 promotes serine36 phosphorylation and mitochondrial translocation of p66Shc

In response to pro-oxidant stimuli, p66Shc becomes phosphorylated at serine36 (S36) and translocates to mitochondria. Once there it oxidizes cytochrome C to produce ROS ^21, 22^, which is a central mechanism to activate the oxidative function of p66Shc. Since the K81 and S36 residues of p66Shc are both present in its conserved CH2 domain, we asked whether promoting SUMO2ylation also facilitates phosphorylation of the S36 residue of p66Shc. In HEK-293 cells as well as in HUVECs, we noted increased levels of p66ShcS36 phosphorylation when SUMO2 and p66Shc were co-expressed (Fig. 4A-D). To determine whether SUMO2-induced S36 phosphorylation is mediated via K81 sumoylation, we evaluated p66ShcS36 phosphorylation in HUVECs overexpressing p66ShcK81R and SUMO2. SUMO2 overexpression significantly increased S36 phosphorylation in cells expressing p66ShcWT whereas this response was markedly attenuated in cells expressing p66ShcK81R (Fig. 4E and F), indicating that K81 SUMOylation promotes, but is not strictly required for, S36 phosphorylation. As p66ShcS36 phosphorylation increases p66Shc level in mitochondria, we evaluated whether SUMO2 affects mitochondrial levels of p66Shc. Levels of p66Shc in the mitochondria of HUVECs increased in a dose-dependent manner upon SUMO2 overexpression (Fig. 5A and B). However, mitochondrial expression of p66Shc was significantly lower in HUVECs expressing p66ShcK81R relative to p66ShcWT (Fig. 5C and D). Collectively, these findings support a model in which SUMO2ylation at K81 facilitates S36 phosphorylation and enhances mitochondrial translocation of p66Shc, thereby promoting its oxidative signaling function.

**Figure 4:**
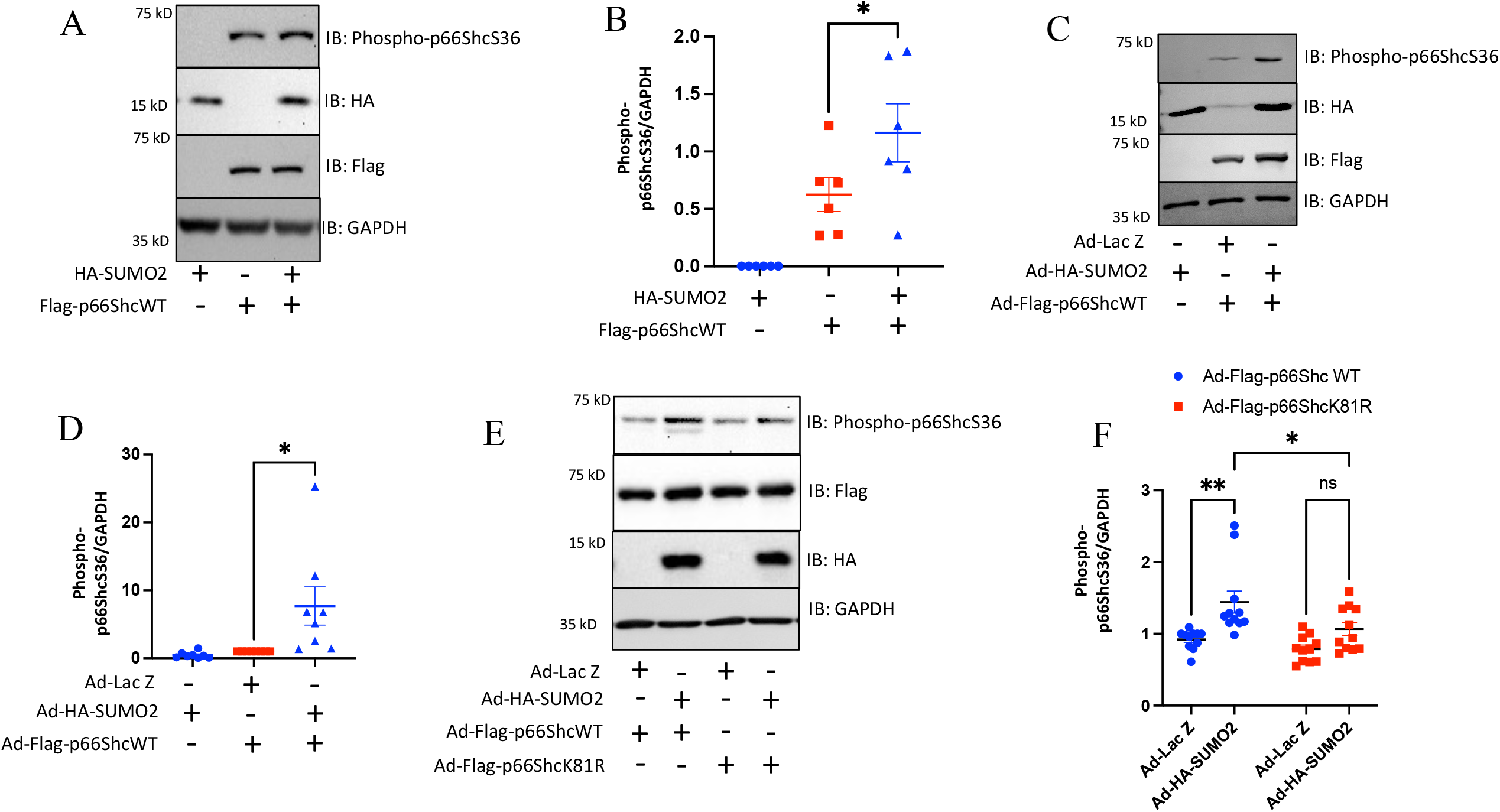
SUMO2 promotes p66ShcS36 phosphorylation by sumoylation of K81. (A) Immunoblot for p66ShcS36 phosphorylation in HEK-293 cells overexpressing SUMO2 (HA-SUMO2) and p66ShcWT (Flag-p66ShcWT). (B) Quantification of p66ShcS36 phosphorylation from A. *P<0.05; n=6; one-way ANOVA. (C) Immunoblot for p66ShcS36 phosphorylation in HUVECs overexpressing SUMO2 (Ad-HA-SUMO2) or p66ShcWT (Ad-Flag-p66ShcWT). (D) Quantification of p66ShcS36 phosphorylation from C. *P<0.05; n=8; one-way ANOVA. (E) Immunoblot for p66ShcS36 phosphorylation in HUVECs expressing SUMO2 (Ad-HA-SUMO2) and p66ShcWT (Ad-Flag-p66ShcWT) or p66ShcK81R (Ad-Flag-p66ShcK81R). (F) Quantification of p66ShcS36 phosphorylation from E. *P<0.05, **P<0.01; n=11; two-way ANOVA. Immunoblots were quantified with iBright image analysis software. Data represents mean ±SEM. Immunoblots are representative of at least three independent experiments.

**Figure 5:**
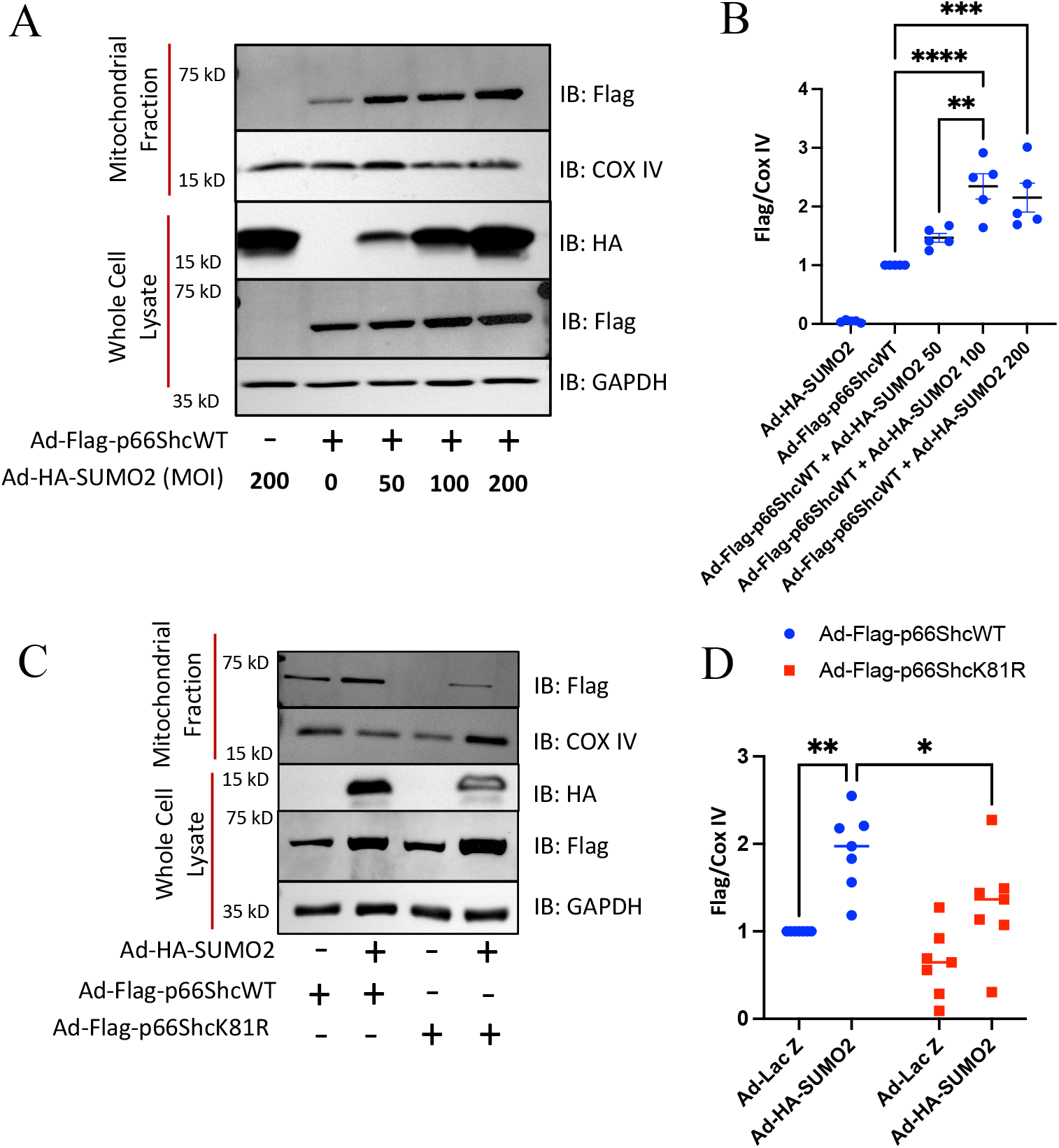
p66ShcK81 SUMO2ylation of K81 in p66Shc promotes its translocation to the mitochondria. (A) Immunoblot for p66ShcWT in whole cell lysates or the mitochondrial fraction of HUVECs ectopically expressing SUMO2 (Ad-HA-SUMO2) and p66ShcWT (Ad-Flag-p66ShcWT). (B) Quantification of the relative abundance of p66Shc in the mitochondria from A. **P<0.01, ****P<0.0001; n=5; one-way ANOVA. (C) Immunoblot for p66Shc in whole cell lysates or the mitochondrial fraction of HUVECs ectopically expressing SUMO2 (Ad-HA-SUMO2) and p66ShcWT (Ad-Flag-p66ShcWT or p66ShcK81R (Ad-Flag-p66ShcK81R). (D) Quantification of relative abundance of p66Shc in the mitochondria from C. *P<0.05, **P<0.01; n=7; two-way ANOVA. Data represents mean ±SEM. Immunoblots are representative of at least three independent experiments.

### Mice expressing non-sumoylatable p66ShcK81R are protected against SUMO2-induced ED

To determine whether SUMO2ylation of p66Shc at lysine 81 modulates endothelial control of vascular tone, we generated a knock-in mouse model in which endogenous p66Shc was replaced with a SUMOylation-deficient mutant (p66ShcK81R). This model was generated using CRISPR-Cas9-mediated genome editing to substitute arginine for lysine at position 81 of p66Shc (Fig. 6A and B). These mice are viable and there was no obvious physiological difference from wild type mice, except that the endothelium-independent relaxation of aortic rings was more pronounced in p66ShcK81R knockin mice (Fig. S4). To examine the effect of SUMO2 on endothelial function in these mice, we used adenoviruses to express SUMO2 in aortic rings from WT and p66ShcK81R knockin mice (Fig. 6C and D). Subsequently, the aortic rings were evaluated for endothelium-dependent relaxation (induced by acetylcholine) and endothelium-independent relaxation (induced by sodium nitroprusside). In aortic rings from WT mice, endothelium-dependent relaxation was significantly delayed by SUMO2 overexpression (Fig. 6E) whereas it was unchanged in p66ShcK81R mice (Fig. 6F). In contrast, endothelium-independent relaxation was not affected by SUMO2 overexpression (Fig. 6G and H). This suggests that the effect of SUMO2 on endothelial function is primarily mediated via p66ShcK81.

**Figure 6:**
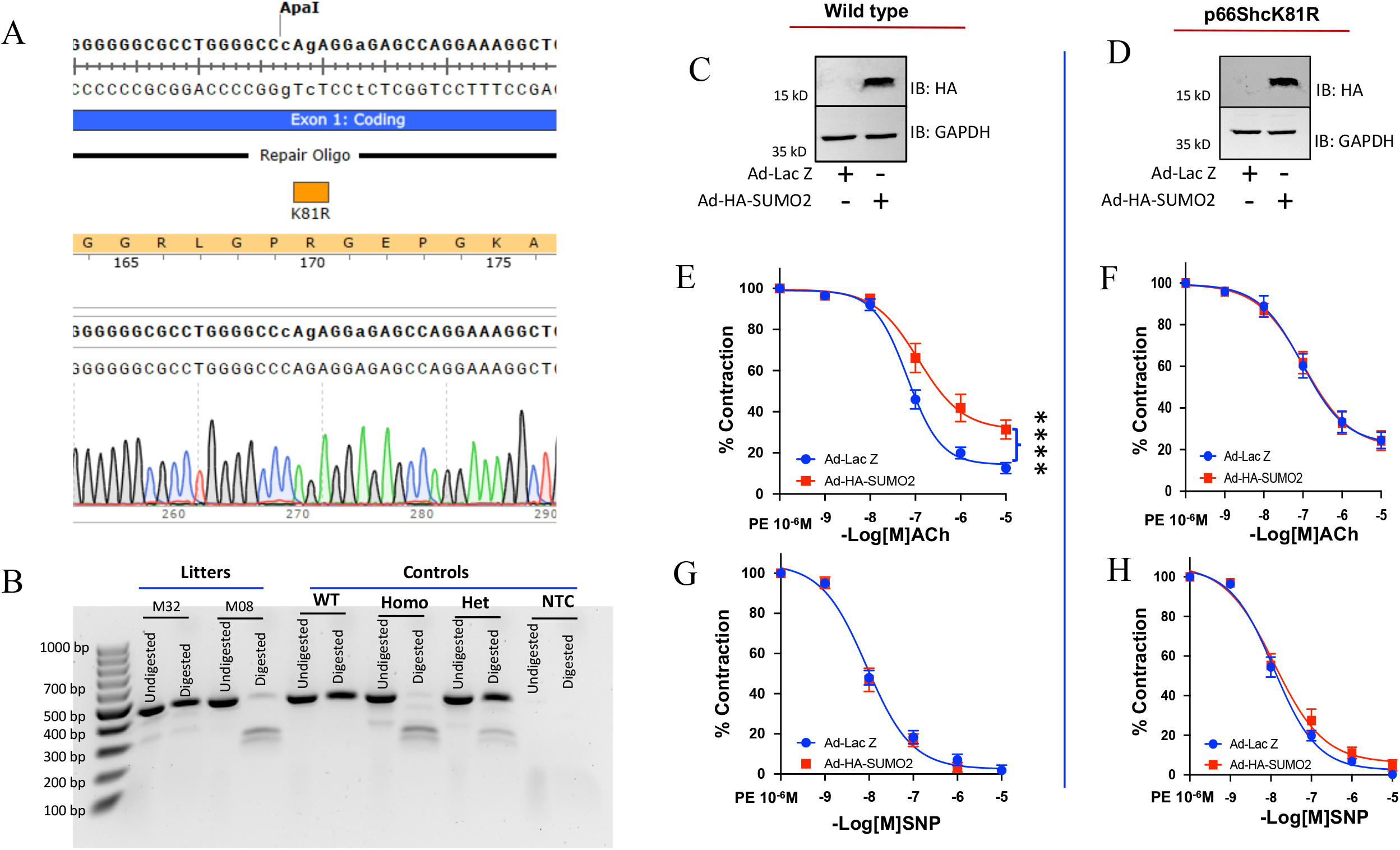
p66ShcK81R knockin (KI) are resistant to SUMO2-induced endothelial dysfunction. (A) DNA sequencing data from a p66ShcK81R knockin (KI) mouse showing a point mutation (A>G) resulting in an amino acid change (K→R), and insertion of silent mutation to create a digestion site for ApaI endonuclease for genotyping. (B) DNA gel electrophoresis of a PCR-amplified and ApaI-digested amplicon identifying the p66ShcK81R KI allele. (C and D) Representative immunoblots showing SUMO2 expression in aortic rings from wild type and p66ShcK81R KI mice (1.67 × 10^8^ pfu/ring). (E and F) Graphs showing % contraction following induction of endothelium-dependent relaxation in aortic rings from (E) wild-type mice, n=14-16 from 4 mice) and (F) p66ShcK81R KI mice (n=15-16 rings from 4 mice) expressing a control vector (Ad-Lac Z) or overexpressing SUMO2 (Ad-HA-SUMO2). (G and H) Graphs showing % contraction following induction of endothelium-independent relaxation in aortic rings from (G) wild type (n=14-16 from 4 mice) and (H) p66ShcK81R KI mice (n=14-15 from 4 mice) expressing a control vector (Ad-Lac Z) or overexpressing SUMO2 (Ad-HA-SUMO2). Curves were compared for nonlinear fit. ****P<0.0001. Ach-acetylcholine, SNP-sodium nitroprusside, PE-Phenylephrine, Homo-homozygous, Het-heterozygous, and NTC-No template control. Data represents mean ±SEM.

### P66ShcK81R knockin mice are protected against hyperlipidemia-induced endothelial dysfunction

To examine the physiological significance of p66Shc-SUMO2, we chose hyperlipidemia as a suitable stimulus as it has been shown to promote sumoylation.^23^ To confirm hyperlipidemia-induced activation of sumoylation and involvement of SUMO2 specifically, we knocked down SUMO2 in HUVECs and treated them with oxidized low-density lipoprotein (o-LDL). Immunoblotting with anti-SUMO2/3 antibody showed a robust increase in sumoylation with o-LDL and the majority of sumoylation was due to SUMO2 (Fig. 7A). Next, we crossbred p66ShcK81R knockin mice with low-density lipoprotein receptor knockout (LDLr^-/-^) mice to generate LDLr^-/-^Xp66ShcK81R KI mice and fed them high fat diet (HFD) for 4 weeks to induce hyperlipidemia. There was no difference in serum cholesterol level between the LDLr^-/-^ and LDLr^-/-^Xp66ShcK81R KI mice on HFD (Fig. 7B). We performed vascular reactivity assay on the aortic rings of WT mice, LDLr^-/-^ mice on normal diet and LDLr^-/-^ on HFD and noted to a significant impairment of endothelium-dependent relaxation of aortic rings from LDLr^-/-^ mice compared to aortic rings from wild type mice and LDLr^-/-^Xp66ShcK81R KI (Fig. 7C). However, the endothelium-independent relaxations were similar among the groups (Fig. 7D). Further we evaluated the level of oxidative stress in aortas of LDLr^-/-^ and LDLr^-/-^Xp66ShcK81R KI mice on HFD by determiening the level of 8-OHdG (8-hydroxy deoxyguansine), a marker of oxidative damage of DNA. We noticed that LDLr^-/-^Xp66ShcK81R KI mice have significantly less oxidative damage to aortic endothelium compared to LDLr^-/-^ mice (Fig. 7E and F). Collectively, our study shows that p66Shc mediates the effects of SUMO2 on vasculature.

**Figure 7.**
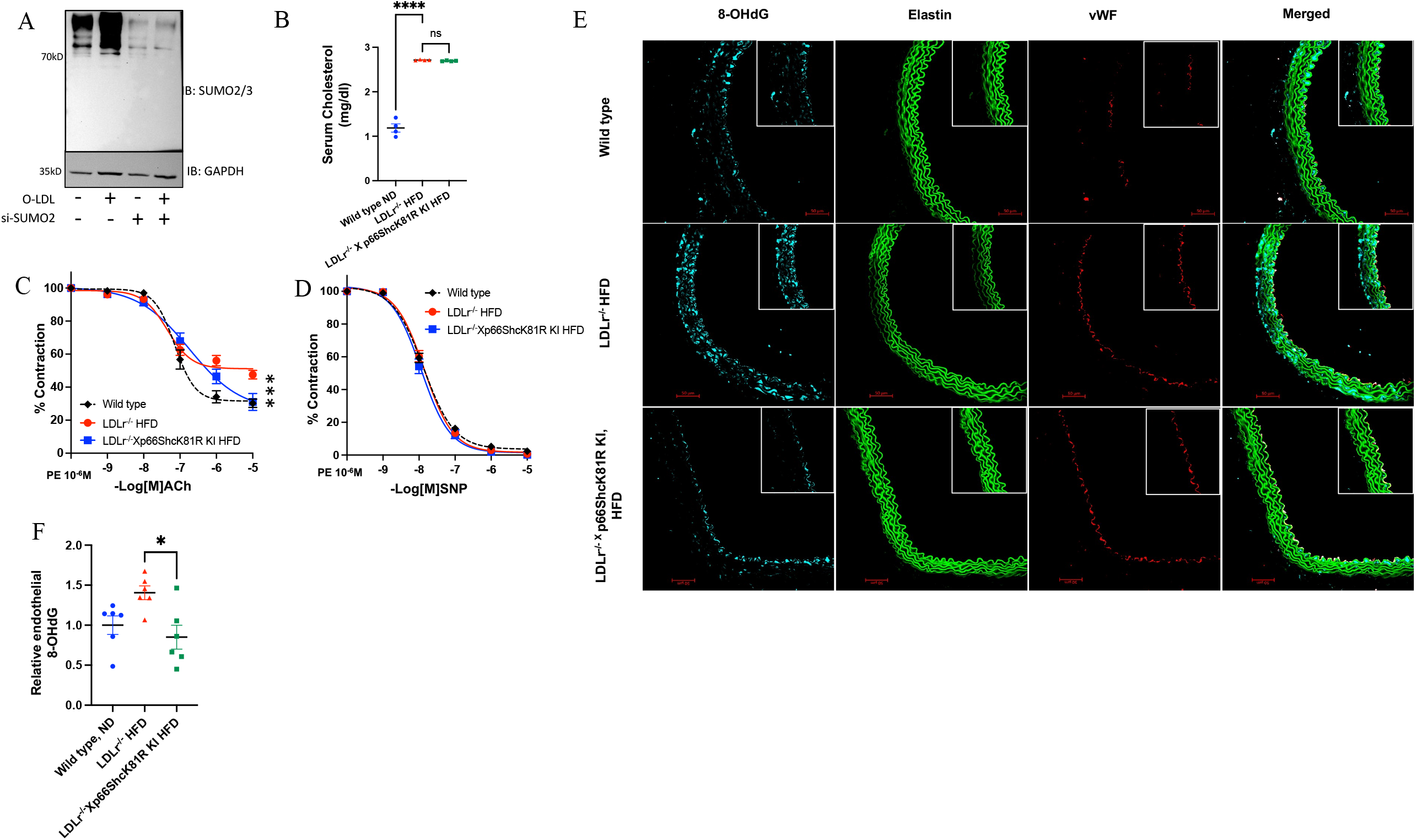
p66ShcK81R knockin (KI) prevents hyperlipidemia-induced endothelial dysfunction and oxidative stress. (A) Immunoblot showing oxidized-LDL (o-LDL)-induced changes in SUMO2/3-conjugated proteins in HUVECs with and without SUMO2. (B) Graph showing serum cholesterol-level in wild type, LDLr^-/-^ mice on high fat diet, and p66ShcK81R KI mice on high fat diet (n=4/group). One-way ANOVA, ****P<0.0001. Graphs showing % contraction following induction of endothelium-dependent relaxation (C) and endothelium-independent relaxation (D) in aortic rings from male wild-type mice, (n=14 rings from 5 mice), LDLr^-/-^ mice on high fat diet (n=18 rings from 6 mice) and LDLr^-/-^ Xp66ShcK81R KI mice on high fat diet (n=12 rings from 6 mice). Two-way ANOVA. ***P<0.001. (E) Representative images showing the level of 8-OHdG in aortic section of LDLr^-/-^ mice on high fat diet (n=6 mice) and LDLr^-/-^ Xp66ShcK81R KI mice on high fat diet (n=6 mice) and their (F) quantification. One-way ANOVA. *P<0.05. HFD-high fat diet Ach-acetylcholine, SNP-sodium nitroprusside, PE-Phenylephrine, 8-OHdG-8 hydroxy deoxyguanosine. Data represents mean ±SEM.

### p66ShcK81-SUMO2ylation affects multiple signaling pathways in endothelial cells

Given that SUMO2 modifies p66Shc at K81 and causes endothelial dysfunction, we next asked how p66ShcK81-SUMO2 affects endothelial cells. We performed global proteomics profiling of HUVECs overexpressing SUMO2 and p66ShcWT or p66ShcK81R using mass spectrometry. Comparative analysis of protein expression was performed against control (HUVECs infected with an equal number of adenovirsues expressing LacZ) (Fig. S5A). Principal component analysis showed a distinct protein expression profile for cells expressing p66ShcWT and p66ShcK81R (Fig. 8A and B). To understand the effect of p66Shc K81 SUMOylation, we examined pathways altered in cells expressing p66ShcWT and p66ShcK81R. The majority of pathways were downregulated, with particularly strong suppression of JAK–STAT signaling. This observation is consistent with the known role of Shc proteins in mediating growth factor receptor signaling. Unlike p52Shc and p46Shc, p66Shc is reported to negatively regulate growth receptor signaling. Supporting this, our mass spectrometry data revealed a greater than 15-fold downregulation of JAK-STAT signaling in cells expressing p66ShcWT in the presence of SUMO2 (Fig. 8C and D). In contrast, this effect was attenuated in cells expressing p66ShcK81R with SUMO2. It is important to note that endogenous p66ShcWT was present in both conditions, which may have competed with the overexpressed p66ShcK81R. Together, these findings suggest that K81 SUMOylation plays a critical role in regulating p66Shc function.

**Figure 8.**
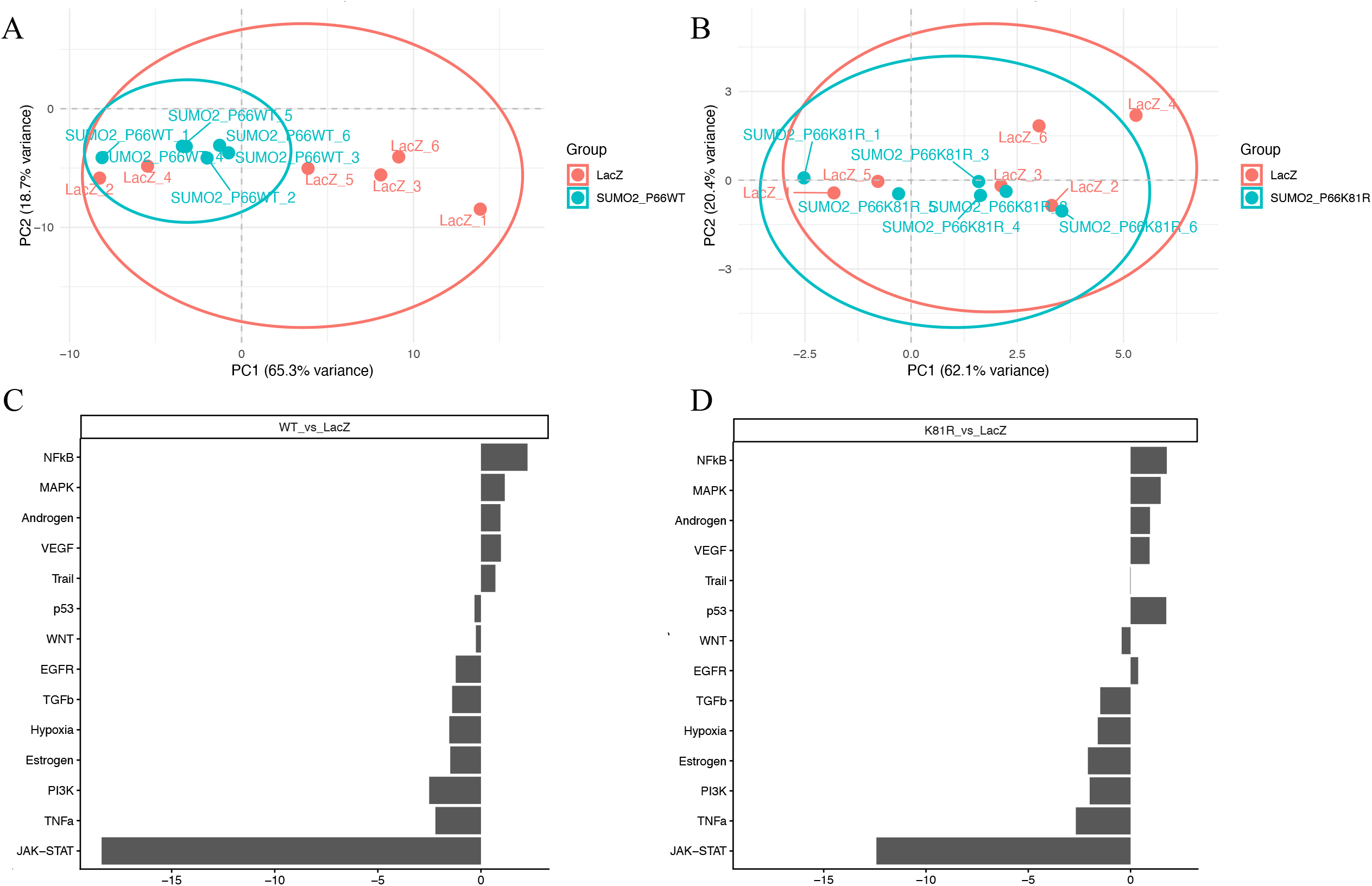
p66ShcK81 sumoylation affects multiple key regulatory pathways in endothelial cells. (A&B) Principal component analysis (PCA) of proteins expressed in endothelial cells expressing SUMO2+p66ShcWT and SUMO2+p66ShcK81R. (C&D) Bar chart presentation of key regulatory pathways differentially affected by SUMO2-p66ShcWT (C) and SUMO2-p66ShcK81R (D) in endothelial cells. The protein expression data (fold change) was obtained from HUVECs expressing SUMO2 with p66ShcWT or p66ShcK81R compared to those expressing Ad-Lac Z. Pathway enrichment analysis was performed using R-based bioinformatics tools, including PROGENy/decoupleR pathway activity inference and visualization packages. Data were derived from a sample size of n=6.

## Discussion

Although SUMO2 and p66Shc are expressed in endothelium and are associated with ED, the mechanisms by which this occurs are unclear. Here we identified SUMO2 modification of p66Shc as a novel molecular signal regulating vascular endothelial function. SUMO2 modifies p66Shc at K81, which increases phosphorylation of p66Shc at S36, resulting in its translocation to the mitochondria. These steps are critical for p66Shc-dependent ROS production. Notably, this phenomenon was evident in non-endothelial cells as well, suggesting that SUMO2-p66Shc may also regulate other physiologic functions. Mice expressing a non-sumoylatable form of p66Shc (p66ShcK81R KI) were protected against SUMO2 and hyperlipidemia-induced ED and oxidative stress, which further supports the role of SUMO2-p66ShcK81 in regulating endothelium function.

Multiple studies suggest that sumoylation promotes ED.^24^ Preventing sumoylation by downregulating the SUMO ligases, Ubc9 (E2 ligase) and PIAS1 (E3 ligase), rescued oxidative stress and advanced glycation end product-mediated endothelial inflammation and dysfunction.^25^ Conversely, partial genetic deletion of the de-sumoylation enzyme, sentrin/SUMO-specific protease 2 (SENP2), significantly increased endothelial apoptosis, inflammation, and dysfunction, which promoted atherosclerotic plaque formation.^5, 6^ However, the isoform-specific effects of SUMOs on endothelial function have not been well studied. We previously reported that endothelial expression of SUMO2 impairs vasorelaxation.^11^ By showing that mice having p66ShcK81R knockin are resistant to endothelial effects of SUMO2 and hyperlipidemia, this study explains the prior findings that the effects of SUMO2 on vasculature are likely mediated by p66Shc.

The adapter protein p66Shc mediates receptor tyrosine kinase signaling in a variety of cell types. Among them, the effects of p66Shc on endothelial cells are well studied and pronounced.^18^ In particular, endothelial function in mice with genetic deletion of p66Shc are protected against a variety of stimuli such as diabetes, aging, and hyperlipidemia.^19, 20, 26^ p66Shc produces ROS by oxidation of cytochrome C in the mitochondria as well as by activating Rac1.^13, 22^ Phosphorylation at S36 in the CH2 domain of p66Shc is one of the earliest discovered posttranslational modifications that govern its oxidative function.^12^ Crosstalk among different posttranslational modifications including phosphorylation, acetylation, and sumoylation is well known.^27-30^ Our finding that SUMO2 increases S36 phosphorylation in p66Shc suggests the existence of such a crosstalk in regulating the oxidative function of p66Shc. The molecular weight of the phosphorylated p66ShcS36 protein band (close to 70kD) suggests that the pool of SUMO2-induced phosphorylated p66ShcS36 is not SUMO2ylated. Therefore, it is likely that K81-SUMOylated and S36-phosphorylated forms of p66Shc coexist in the same pool of protein, but that p66Shc is not SUMOylated and phosphorylated at the same time. Future studies will be needed to establish the hierarchy of this phenomena.

Based on mass spectrometry analysis and *in vitro* sumoylation of recombinant p66Shc, it appears that p66ShcK81 is a primary target of SUMO2. However, the multiple protein bands higher than 66kD that were present in the immunoprecipitation assay suggest the possibility that additional sumoylation sites are present in p66Shc. Technical challenges limited our ability to determine the nature of these protein bands. Yet, in silico analysis using an available sumoylation prediction tool (Cuckoo workgroup) indicates 3 possible sumoylation motifs present in p66Shc, namely K81, K462, and K582, which can non-specifically be modified by SUMO1/2/3/4. Importantly, among the possible sumoylation motifs only K81 is present in the CH2 domain, which is unique structural feature of p66Shc (i.e., not present in other isoforms: p52Shc and p46Shc) that has a regulatory effect on its function. Therefore, even though other lysines are sumoylated, it is likely that the effect of p66Shc-sumoylation is mediated through the K81 modification and has also been supported by proteomics analysis.

The primary function of the ShcA family of adaptor proteins is to mediate growth factor receptor signaling ^18^. However, p66Shc is unique within this family, as it deviates from the classical role of Shc proteins and instead promotes oxidative stress. Our global mass spectrometry data suggest that enhancing p66ShcK81 SUMO2ylation inhibits JAK-STAT signaling, a key downstream pathway of growth factor receptor signaling. The negative regulation of growth receptor signaling by p66Shc, through competition with p52Shc for Grb2 binding, has been previously reported ^31^. In addition, a recent study demonstrated that p66Shc can mediate EGFR-ERK signaling ^32^. Therefore, beyond its established role in ROS regulation, p66ShcK81-SUMO2ylation may have additional functional roles in endothelial cell physiology.

One of the limitations of this study is that we did not perform gain-of-function experiments to ascertain if SUMO2-modified p66ShcK81 regulates p66Shc function *in vivo*, which is technically too challenging. Additionally, we were unable to examine the behavior of the phosphomimetic double mutant (S36D/K81R) to directly assess the interplay between K81 SUMO2ylation and S36 phosphorylation in mitochondrial localization. Prior studies have shown that K81 of p66Shc undergoes acetylation^14^ and that a specific lysine can undergo sumoylation as well as acetylation.^29^ Although p66ShcK81R disrupts acetylation and sumoylation that occurs from acetyl transferase and SUMOylating enzymes, it is possible that the same lysine is sumoylated and acetylated in a context dependent manner. Importantly, the extent of post translational modification is so small that a competition between acetylation and sumoylation for the same target residue is highly unlikely. The fact that p66ShcK81R KI mice are protected against SUMO2-induced ED supports the hypothesis that p66ShcK81 mediates the endothelial effects of SUMO2.

Collectively, our study identified p66Shc SUMO2ylation as a unique cellular signaling mechanism specifically regulating vascular endothelial function. As such, this signaling mechanism may also affect advanced pathologies such as atherosclerosis, where both increased SUMOylation and expression of p66Shc has been reported. Nevertheless, given its potential to regulate endothelial function, SUMO2ylation of p66Shc presents a novel therapeutic target for pathologies associated with ED.

## Acknowledgements

S. Kumar was supported by NIH grants R01HL152132, R21AG086743 and CFF pilot and feasibility grant 005803I223. Sequencing data presented herein were obtained at the Genomics Division of the Iowa Institute of Human Genetics, which is supported, in part, by the University of Iowa Carver College of Medicine. Viral Vectors were provided by the University of Iowa Viral Vector Core. Mass spectrometry was performed at Mass Spectrometry and Proteomics Core Facility, University of Nebraska. The University of Nebraska Medical Center Mass Spectrometry and Proteomics Core Facility is administrated through the Office of the Vice Chancellor for Research and supported by state funds from the Nebraska Research Initiative. The p66ShcK81R knockin mouse was generated at the University of Iowa Genome Editing Core Facility, supported in part by grants from the NIH and from the University of Iowa Carver College of Medicine. We wish to thank Norma Sinclair, Patricia Yarolem, Joanne Schwarting and Rongbin Guan for their technical expertise in generating transgenic mice. We gratefully acknowledge Kris Griener of the Design Center for language editing and Dr. Jennifer Y. Barr of the Scientific Editing and Research Communication Core at the University of Iowa Carver College of Medicine for critical reading of the manuscript and suggestions toward improving the clarity of the manuscript.

## Notes

**Competing Interest Statement:** The authors have declared that no conflicts of interest exist.

### Competing Interest Statement

The authors have declared no competing interest.

### Summary of Updates

The manuscript has been reviewed by external reviewers and has been modified in accordance with their suggestions.

