## Supplement file for "p66Shc Mediates SUMO2-induced Endothelial Dysfunction"

**Materials and Methods**

**Animals:** All experimental animal procedures were approved by the Institutional Animal Care and Use Committee at the University of Iowa and conformed to the Guide for the Care and Use of Laboratory Animals published by the National Institutes of Health. Wild-type mice (C57BL/6) were procured from Jackson lab. The p66ShcK81R knockin mice were generated with the help of Genome Editing Core Facility, University of Iowa, Iowa City. Using a CRISPR-Cas9 technique, a mutation was introduced in the embryos from C57BL/6J mice and the embryos were implanted to pseudo-pregnant females. Founders were identified by PCR genotyping and Sanger sequencing. Subsequently mice were inbred to generate homozygous p66ShcK81R knockin mice.

To generate LDLr^-/-^/p66ShcK81R KI mice, LDLr^-/-^ mice (B6.129S7-*Ldlr^tm1Her^*/J) were obtained from Jackson lab and crossbred to p66ShcK81R KI mice to generate LDLr^-/-^/p66ShcK81R KI mice homozygous for both LDLr deletion and p66ShcK81R knockin. To induce hyperlipidemia, these mice were fed on high fat diet (Envigo-TD.88317) for 4 weeks. Serum cholesterol was measured using a commercially available kit (Cell Biolab, #STA-390).

**Genotyping protocol for p66ShcK81R**: DNA was isolated from ear punches of mice using a commercially available DNA isolation kit (NucleoSpin Tissue DNA, Machery-Nagel GmbH & Co.). Polymerase chain amplification (PCR) was performed to amplify the DNA sequence around the knockin position using Phusion high-fidelity DNA polymerase master mixture reaction (NEB); forward primer (5’- CTT CGG AAT GAG TCT CTG TCA TC-3’) and reverse primer (5’- ACC CGA ACC AAG TAG GAA ACG-3’). The reaction mixture was pre-heated at 70°C for 2 min followed by initial denaturation at 98°C for 2 min, then 35 cycles at 98°C for 20s, 62°C for 30s, 72°C for 40 sec, and a final extension at 72°C for 5min. PCR amplified products were digested with ApaI enzyme at 37°C for 2.5 h. Agarose gel electrophoresis was performed to determine the amplification and digestion pattern of the amplicon. Mice identified by genome sequencing were used as a positive control.

**Cell culture:** Human umbilical vein endothelial cells (HUVECs) were purchased from Promocell (Heidelberg, Germany) and Lonza (Walkersville, MD) and were cultured in endothelial growth medium-2 from Promocell and Lonza. HEK 293 cells were cultured in DMEM (Gibco) supplemented with 10% fetal bovine serum (Thermo), and 1% Pen Strep (penicillin 10,000 Units/mL and 10,000 ug/mL streptomycin; Gibco).

**Reagents:** Primary antibodies: Monoclonal Flag (Catalog no. F1804, Sigma, MO); Monoclonal GAPDH (Catalog no. 2118S, Cell Signaling Technology, MA); monoclonal Senp1 (Catalog no. sc- 271360, Santa Cruz Biotechnology, TX); Monoclonal HA (Catalog no. 3724s Cell Signaling Technology); Monoclonal His-tag (Catalog no. 2366S Cell Signaling Technology); Polyclonal Phospho-p66Shc (Catalog no. Ab267413 Abcam, MA); SUMO2/3 (Catalog no. 633306, Sigma and Catalog no. 4971, Cell Signaling Technologies); Monoclonal Ubc9 (D26F2) (Catalog no. XP 4786s, Cell Signaling Technology); PhosphoDetect Anti-Shc/p66 (Catalog no. 566807, Millipore Sigma, MO); Cox IV (3E11) (Catalog no. 4850S, Cell Signaling Technology)**;** sumoylation kit (BML-UW8955-0001, Enzo Life Sciences, NY).

Secondary antibodies: polyclonal IR-dye 800 Rb (Catalog no. ab216773, Abcam); polyclonal IR-dye 680 (Catalog no. ab216776, Abcam); anti-rabbit IgG HRP linked (Catalog no. 7074S, Cell Signaling Technology); anti-mouse IgG HRP linked (Catalog no. 7076S, Cell Signaling Technology).

**Plasmids**: We generated a new construct of p66Shc (human) with a Flag tag (DYKDDDDK) at the N-terminus (Flag-p66Shc) in the pcDNA3 backbone. FNpCDNA3 was a gift from Dr. Robert Oshima (Addgene plasmid # 45346). We also generated HA-tagged (MYPYDVPDYA) SUMO2 (Human) in the same backbone. Successful cloning was verified by DNA sequencing. FLAG-SENP1 (Addgene plasmid # 17357) ^1^ and Ubc9 (Addgene plasmid # 20082) ^2^ was a gift from Dr. Edward Yeh.

**Adenoviruses**: Adenoviruses of Flag-p66ShcWT, Flag-p66ShcK81R, and HA-SUMO2 were generated with the help of Viral Vector Core facility, University of Iowa. Target sequences were cloned to a shuttle vector. For adenoviral expression, Flag-p66ShcWT, Flag-p66ShcK81R, and HA-SUMO2 were cloned into a shuttle vector plasmid (G0688pacAd5CMV) provided by the Viral Vector Core facility. Constructs were verified for proper inserts by digestion followed by Sanger sequencing. Further processing was supported by the Viral Vector Core facility.

**Mutagenesis of p66Shc**: To create p66ShcK81R, we performed site-directed mutagenesis on Flag-p66Shc to induce an AAG to AGA mutation. The following primers were used: forward 5'-cctggctcccctctagaccctgggcgcc-3' and reverse 5'-ggcgcccagggtctagaggggagccagg-3'. Successful mutagenesis was verified by DNA sequencing.

**Adenoviral expression:** An optimum volume of viral stock was added to the cells with growth medium. We used 1:100 MOI in most of the experiments. Specific MOI is described in the figures Cells were analyzed almost 40 h of infection.

**Plasmid and siRNA transfection:** For plasmids transfection, Lipofectamine 2000 (Invitrogen, MA) was used as per the manufacturer instructions. For 1 μg of plasmid, we used 2 μl lipofectamine. p66Shc-specific siRNAs (custom made) and a non-specific control siRNA were purchased from Invitrogen. An appropriate amount of siRNA was incubated with Lipofectamine2000 (Invitrogen) at room temperature for 20 min before adding to cells prepared with OPTI-MEM media (Invitrogen). Four hours later the media was replaced with fresh endothelial growth media and cells were cultured for an additional 48 hours.

**Recombinant protein expression and purification:** Purified recombinant protein was synthesized as published earlier with slight modification ^3^. DNA sequences coding His-p66ShcWT and His-p66ShcK81R proteins were PCR-amplified from Flag-p66ShcWT and Flag-p66ShcK81R plasmids. PCR amplified His-p66ShcWT and His-p66ShcK81R were cloned into the pET 21a expression vector. Sanger sequencing confirmed the presence of the insert. Sequenced plasmids were transformed into Rosetta 2 (de3) cells and inoculated into 50 ml of LB broth supplemented with 50 μl ampicillin (100μg/ml) and 50 μl chloramphenicol (34μg/ml) antibiotics. The cells were then incubated overnight at 37°C at 250 rpm for growth. For secondary inoculum, 10 ml of the primary culture was inoculated into 1L of 2xYT media (Yeast-Tryptone) and incubated at 37°C until the O.D. reached 0.6 at 600nm. Cultures were then cooled at 2-8°C for a minimum of 20 minutes, followed by 5 minutes at room temperature. Induction of the culture was done using 0.250 mM IPTG, and the incubation continued at 16°C, 200 rpm overnight. The culture was pelleted down by centrifugation at 4°C. The cell pellets were sonicated on ice (five 30s pulses). The lysed sample was centrifuged at 18000 rpm for one hour at 4°C. The supernatant was collected and passed through a pre-packed, pre-equilibrated Ni-NTA resin column for protein purification. Western blotting confirmed the presence of the target protein.

**Mass spectrometry for sumoylation and Data Analysis:** Recombinant p66Shc was sumoylated with a commercially available sumoylation kit (Enzo Life Sciences). Sumoylated protein was resolved by SDS-PAGE. Gels were stained with mass spectrometry compatible silver stain (SilverQuest, Invitrogen). Gel pieces were excised based on the observed SUMOylation signal in the immunoblot of recombinant p66Shc. Gel pieces were de-stained and reduced with 5 μL of 200 mM tris(2-carboxyethyl) phosphine (TCEP) (1 h incubation, 55 ⁰C) and alkylated with 5 μL of 375 mM iodoacetamide (IAA) (30 min incubation in the dark, room temperature). The protein digestion was carried out using 2.5 μg of trypsin per sample (16 h incubation, 37 ⁰C). The next day, samples were dried out using a speed vacuum and then desalted with C18 spin columns (Pierce, MA). Clean peptides were dried out again with a speed vacuum and resuspended in 0.1% formic acid, and next analyzed using a high-resolution mass spectrometry nano-LC-MS/MS Tribrid system, Orbitrap Fusion™ Lumos™ coupled with UltiMate 3000 HPLC system (Thermo Fisher Scientific). Peptide identification was performed by searching MS/MS data against the p66Shc protein sequence in Peaks Studio. We detected a missed cleavage at a lysine (K) residue as shown in the figure 3B, which was followed by a QTGG modification with a mass of 343.15 Da. This specific mass shift was consistent with the expected tryptic peptide from sumoylated peptide as shown by other studies ^4, 5^.

**Label free quantitative mass spectrometry and data analysis**: HUVECs were infected with Ad-HA-SUMO2 znd Ad-Flag-p66ShcWT or Ad-Flag-p66ShcK81R. Samples were lysed in RIPA buffer and protein expression was determined by immunoblotting. Whole cell lysate having 50 µg of protein per sample from six biological replicates per group was taken and detergent was removed by chloroform/methanol extraction, and the protein pellet was re-suspended in 100 mM ammonium bicarbonate and digested with MS-grade trypsin (Pierce) overnight at 37°C following reduction with 10 mM DTT at 56°C for 30 mins and alkylation using 50 mM iodoacetamide at RT for 25 mins. Peptides were cleaned with PepClean C18 spin columns (Thermo) and were re-suspended in 2% acetonitrile (ACN) and 0.1% formic acid (FA) and 500 ng of each sample was loaded onto trap column Acclaim PepMap 100 75µm x 2 cm C18 LC Columns (Thermo Scientific™) at flow rate of 4 µl/min then separated with a Thermo RSLC Ultimate 3000 (Thermo Scientific™) on a Thermo Easy-Spray PepMap RSLC C18 75 µm x 50 cm C-18 2 µm column (Thermo Scientific™) with a step gradient of 4–25% solvent B (0.1% FA in 80 % ACN) from 10-100 min and 25–45% solvent B for 100–130 min at 300 nL/min and 56°C with a 155 min total run time. Eluted peptides were analyzed by a Thermo Orbitrap Exploris 480 (Thermo Scientific™) mass spectrometer in a data dependent acquisition mode. A survey full scan MS (from m/z 350–1200) was acquired in the Orbitrap with a resolution of 60,000. The Normalized AGC target for MS1 was set as 300% and ion filling time set as 25 ms. The most intense ions with charge state 2-6 were isolated in 3 s cycle and fragmented using HCD fragmentation with 30% normalized collision energy and detected at a mass resolution of 15,000 at 200 m/z. The AGC target for MS/MS was set as 50% and ion filling time set to auto for 30 s with a 10 ppm mass window. Protein identification was performed by searching MS/MS data against the swiss-prot human protein database using the in house PEAKS X + DB search engine. The search was set up for full tryptic peptides with a maximum of two missed cleavage sites. The precursor mass tolerance threshold was set 10 ppm for and maximum fragment mass error was 0.02 Da. The significance threshold of the ion score was calculated based on a false discovery rate of ≤ 1%. Quantitative data analysis was performed using progenesis QI proteomics 4.2 (Nonlinear Dynamics). Statistical analysis was performed using ANOVA and The Benjamini-Hochberg (BH) method was used to adjust p values for multiple-testing caused false discovery rate. The adjusted p ≤ 0.05 was considered as significant.

Principal Component Analysis (PCA) was employed to assess the global molecular differences induced by SUMO2_P66WT and SUMO2_P66K81R expression relative to the LacZ control. In both comparisons, PCA revealed distinct clustering of experimental groups, indicating substantial divergence in molecular profiles. For the SUMO2_P66WT group, PC1 accounted for 65.3% of the total variance, while PC2 explained 18.7%, together capturing 84% of the dataset’s variability. Similarly, in the SUMO2_P66K81R comparison, PC1 and PC2 explained 62.1% and 20.4% of the variance, respectively, totaling 82.5%. In both cases, biological replicates clustered tightly within their respective groups, underscoring the reproducibility of the data. These results demonstrate that both SUMO2_P66WT and SUMO2_P66K81R expression lead to pronounced and consistent alterations in the molecular landscape compared to LacZ, with each variant contributing distinct signatures of change.

**Whole cell ROS assay by spectrophotometry:** HUVECs were plated in clear-bottom black-walled 96 well plates and infected with adenoviruses encoding HA-SUMO2 or control virus encoding LacZ. For ROS detection and quantification, cells were incubated with MitoSOX (2µM, Life Technologies) in EBM2 for 30 min. at 37°C in CO_2_ incubator. Following incubation, cells were washed with pre-warmed EBM2 media and maintained at 37°C in a CO_2_ incubator for another 30 min. MitoSOX fluorescence was measured at 396/610 nm using a plate reader. Each well was measured at 4 different points at the 1 mm distance and the mean was considered as final reading of that well. Values were normalized against the background fluorescence. We excluded the first and last row in each plate.

**Immunoprecipitation:** Protein lysate was pre-cleared using Sepharose beads for 2 hours at 4°C on the rocker. The supernatant was collected and incubated with the specified antibody (1 ug of antibody/mg protein) and Sepharose beads overnight at 4°C. Sepharose beads were washed 8 times with ice cold lysis buffer (1 ml each wash). After that the beads were boiled for 5 min at 95°C and subjected to SDS-PAGE.

**Generation of SUMO2-p66Shc antibody:** The antibody to detect SUMO2-p66Shc was generated by using the branched peptide, mimicking the region of Sumo2 attachment (Sumo2-K81) to p66Shc as an immunogen using commercial source (YenZym, CA). The antobody was purified and tested for its specificity by ELISA and immunoblotting.

**Immunoblotting:** Cells or tissues were washed with PBS and lysed in an appropriate amount of protein lysis buffer (TritonX100 -1%, Tris 7.4 -50mM, EDTA -5mM, NaCl -150mM, and glycerol -5%) containing Protease Inhibitor Cocktail (Roche). We used Phosphatase Inhibitor Cocktail (Millipore) and de-sumoylation inhibitor N-ethylmaleimide (20 mM) in the lysis buffer where appropriate. Cell lysis was performed by repeated freeze thaw followed by sonication for 25 sec (5 sec on, 2 sec off, and 20% amp). Homogenate was centrifuged at 13000g for 20 mins at 4°C, supernatant was used for protein quantification.

We used approximately 30 ug of protein lysates. Cell lysate was mixed with protein loading dye and boiled at 95 ºC for 5 min. Following brief centrifugation, lysates were loaded onto acrylamide gel for SDS-PAGE. Protein was transferred to a nitrocellulose membrane. Membranes were treated with 5% milk or BSA in tris-buffered saline with 0.1% tween20 and were probed with the specified primary antibody and the appropriate peroxidase-conjugated (Abcam) or fluorescent antibody (Abcam and Licor, NE) secondary antibody. Chemiluminescent signal was developed using Super Signal West Femto substrate (Pierce, Rockford, IL). Images were acquired using a signal detection system (iBright 1500FL, Thermo) and quantified with iBright image analysis software.

**Mitochondrial Translocation**: HUVECs were used to determine the mitochondrial localization of p66Shc. Cells were washed with cold 1xPBS buffer and collected in 500ul MSE buffer (5 mM MOPS, 70 mM sucrose, 2 mM EGTA, 220 mM mannitol, pH 7.2). Maintaining the cell suspension at 4ºC, cells were homogenized using a smooth PTFE pestle in glass tube until 85 to 95 % of the cells were ruptured. Unbroken cells and nuclei were removed by centrifugation at 1000g for 5 min. Supernatant was collected and centrifuge at 5000g for 10 min to obtain the mitochondrial and cytosolic fractions. The mitochondrial pellet was further processed for protein lysate, as described under Immunoblotting).

***Ex vivo* adenoviral infections and Vascular reactivity:** Ad-LacZ or Ad-HA-SUMO2 adenoviral stocks at a final concentration of 2.5 × 10^7^ pfu/ml in EGM2 media were added to each aortic ring and incubated at 37°C for 24h. Infected vessels were suspended between two wire stirrups (150 mm) in a four-chamber myograph system (DMT Instruments) in 6 ml Krebs-Ringer (95% O2-5% CO_2_, pH 7.4, 37°C). All concentration-effect curves were performed on arterial rings beginning at their optimum resting tone. Endothelium-dependent and –independent relaxation was determined by generating dose-response curves to acetylcholine (ACh 10^-9^-10^-5^ M) and sodium nitroprusside (SNP 10^-9^-10^-5^ M), respectively on phenylephrine (PE 10^-6^ M)-induced pre-contracted vessels. Vasorelaxation evoked by ACh and SNP was expressed as percent contraction with reference to phenylephrine (PE 10^-6^ M) and determined by calculating the percentage of inhibition to the pre-constricted tension.

**Histological processing and quantification:** For immunohistochemical examinations 10 μm thick sections were prepared from cryopreserved aortic rings. Sectioned were washed with PBS (3 times) and were blocked with normal goat serum (Millipore) for 30 minutes at room tempereature. Next, the blocking medium was removed, primary antibody was added and sections were incubated at 4°C. After incubation with primary antibodies the sections were incubated with secondary antibodies tagged with fluorescent dye were used. Images were acquired on confocal microscope (Zeiss 710). Images were quantified using Zen Blue software (Zeiss). For endothelial specific measurements, endothelial layer was manually traced following the lead of inner most elastic lamina and staining for endothelial-specific von Willebrand factor. Multiple images were captured from each section and the mean fluorescence intensity was calculated for each image which was averaged for each mouse, and expressed as fold differences among groups. Autofluorescence of elastin lamina was captured to clearly define the endothelial specific fluorescence.

**Statistics:** All data met assumptions of the statistical test, and all statistical analyses were performed using GraphPad Prism 9 unless specified. Data represent the mean ± SEM of at least three independent assays. Normal distribution was checked using Kolmogorov-Smirnov test. Significance of differences between two groups was determined using a two-tailed independent sample Student *t*-test. One-way ANOVA was used for comparing more than two groups and two-way ANOVA was used for comparing two variables simultaneously, followed by Tukey *post-hoc* analysis for multiple group comparison. Vascular reactivity data was analyzed by non-linear regression analysis (curve fit) and significance of differences was determined by comparing two curves for the fitting line. Results were considered significant if P values were ≤ 0.05.

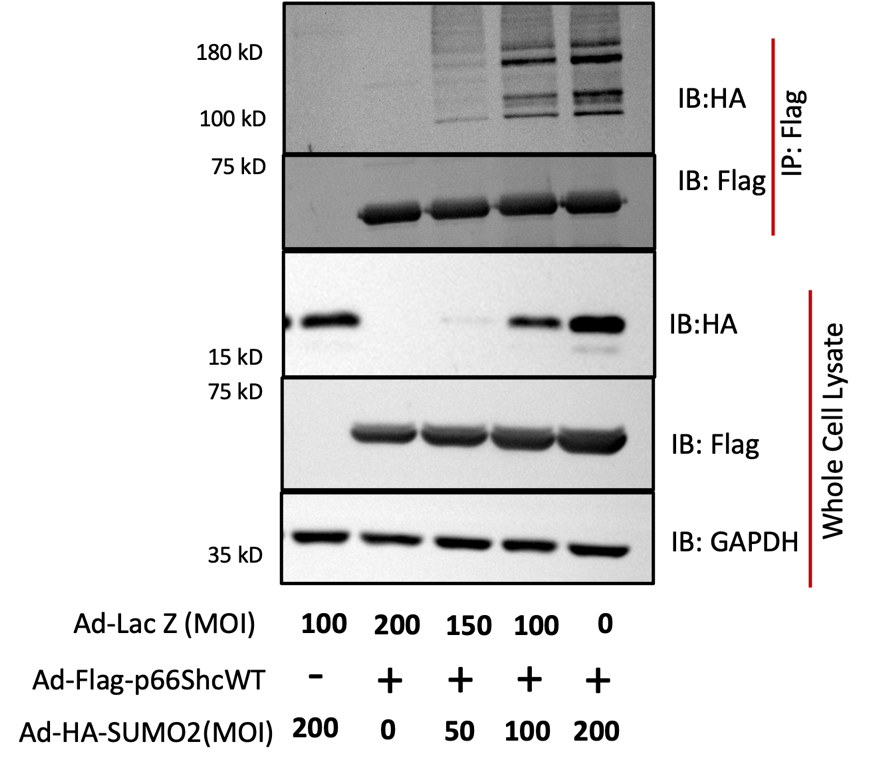

Fig. S1. SUMO2 promotes p66Shc SUMO2ylation in a dose dependent manner. Immunoblot of HUVECs transduced with wild-type p66Shc (Ad-Flag-p66ShcWT) and SUMO2 (Ad-HA-SUMO2) at the indicated concentrations. Immunoblots are representative of at least three independent experiments.

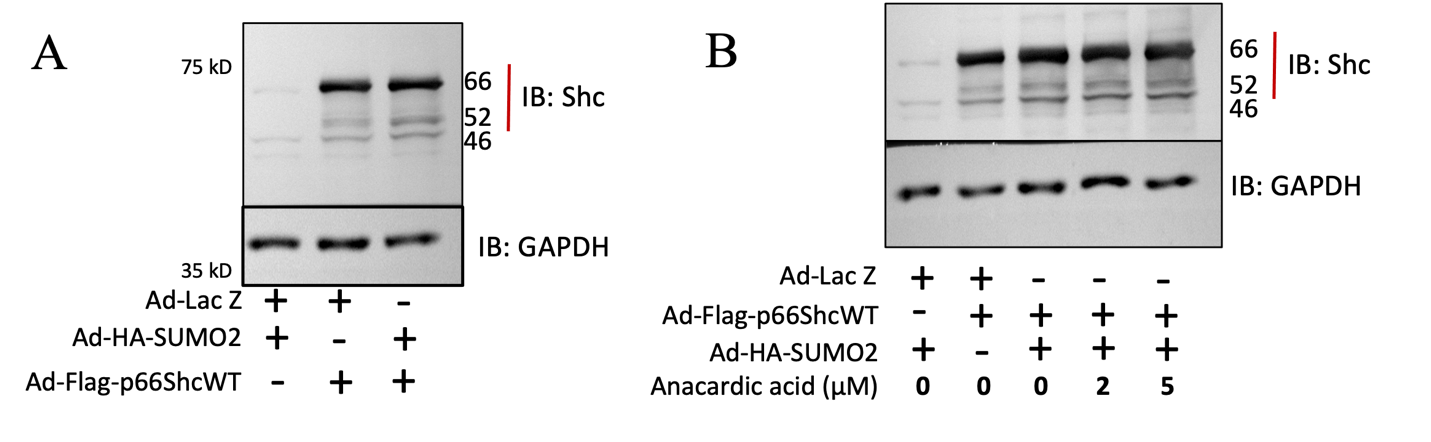

**Figure S2.** Immunoblot showing the relative overexpression level of p66Shc in HUVECs used for (**A**) experiment 2C and (**B**) experiment 2D.

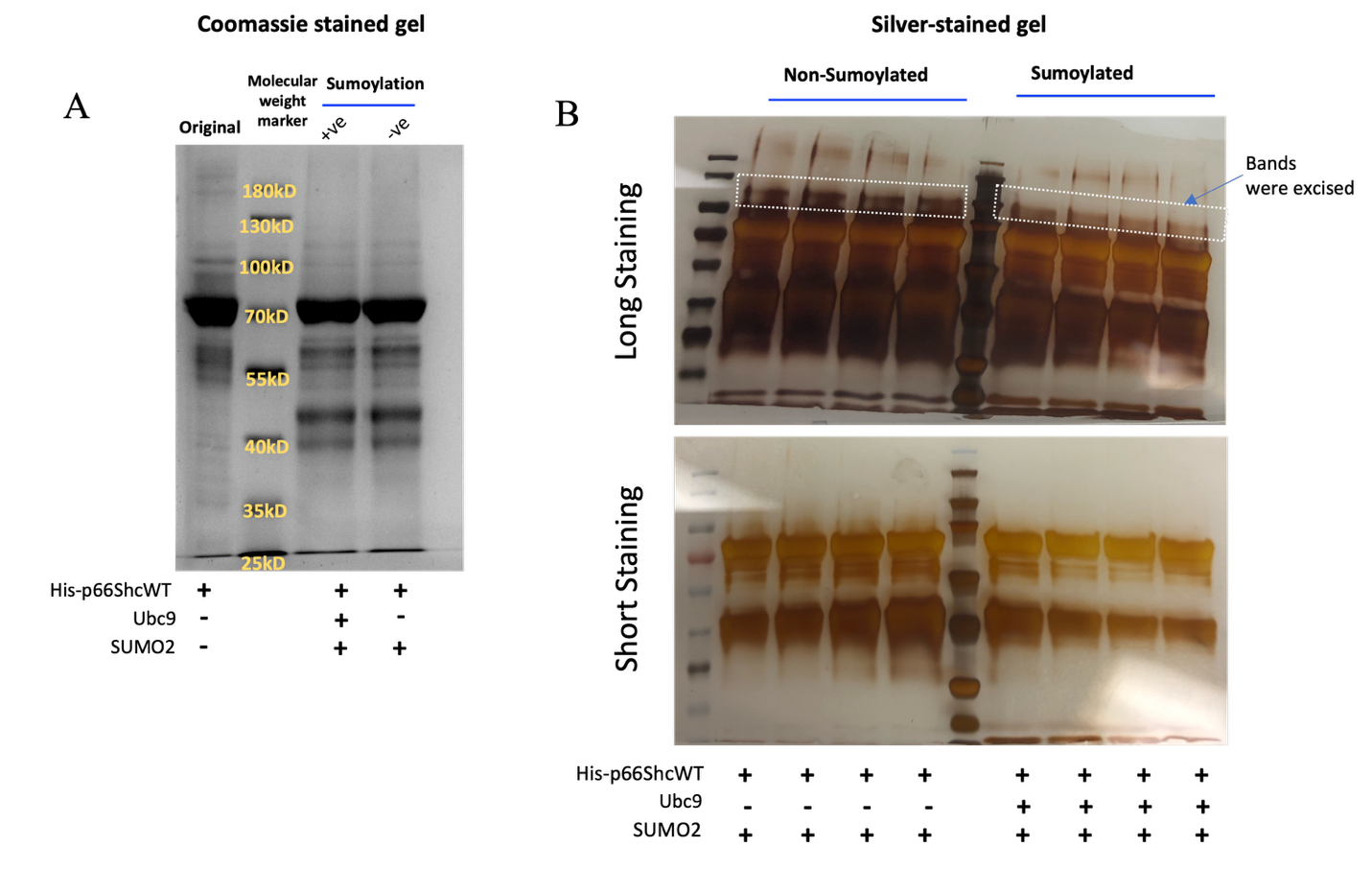

Fig. S3. Sumoylation of recombinant p66ShcWT for mass spectrometry. (A) Coomassie stain of SDS-PAGE of recombinant p66ShcWT in its original form (not processed for SUMO2ylation), and after being SUMO2ylated and non-SUMO2ylated. (B) Silver stain of non-SUMO2ylated and SUMO2ylated recombinant p66ShcWT used for mass spectrometry.

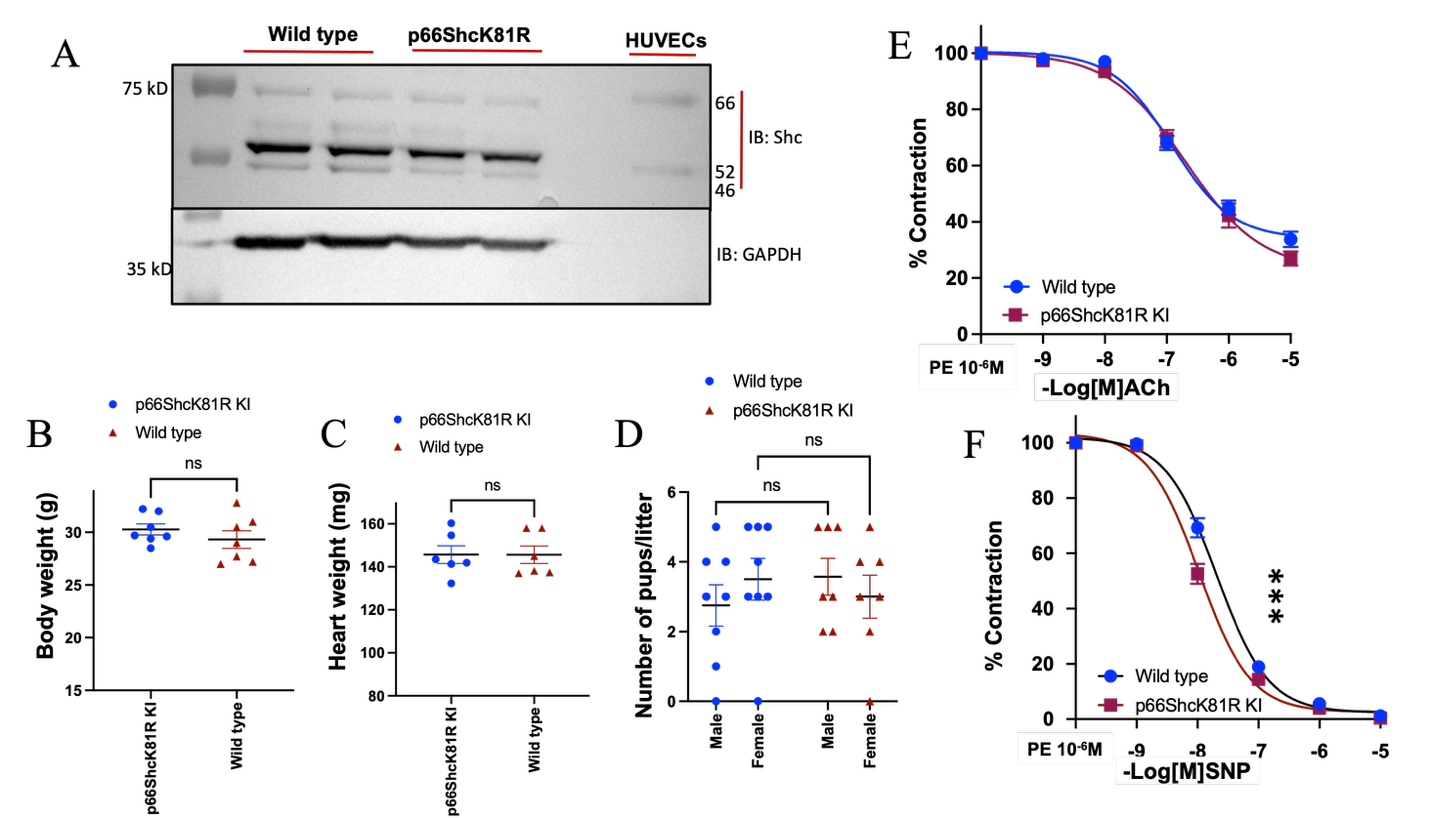

**Fig S4. Physiological characterization of p66ShcK81R KI mice.** (A) Immunoblot for p66Shc isoforms in aortic lysates from wild-type and p66ShcK81R knockin mice. (B) Body weight of p66Shc K81R KI and wild type mice (n=7/group). (C) Heart wight of p66ShcK81R KI and wild type mice (n=6/group). (D) Litter-size of p66ShcK81R KI (n=7) and wild type mice (n=8). Graphs showing % contraction following induction of (E) endothelium-dependent and (F) endothelium-independent relaxation in aortic rings from wild-type mice, (n=13 rings from 4 mice) and p66ShcK81R KI mice (n=14 rings from 6 mice). Curves were compared for nonlinear fit. ***P<0.001. ns= not significant. Ach-acetylcholine, SNP-sodium nitroprusside, PE-Phenylephrine. Data represents mean ±SEM.

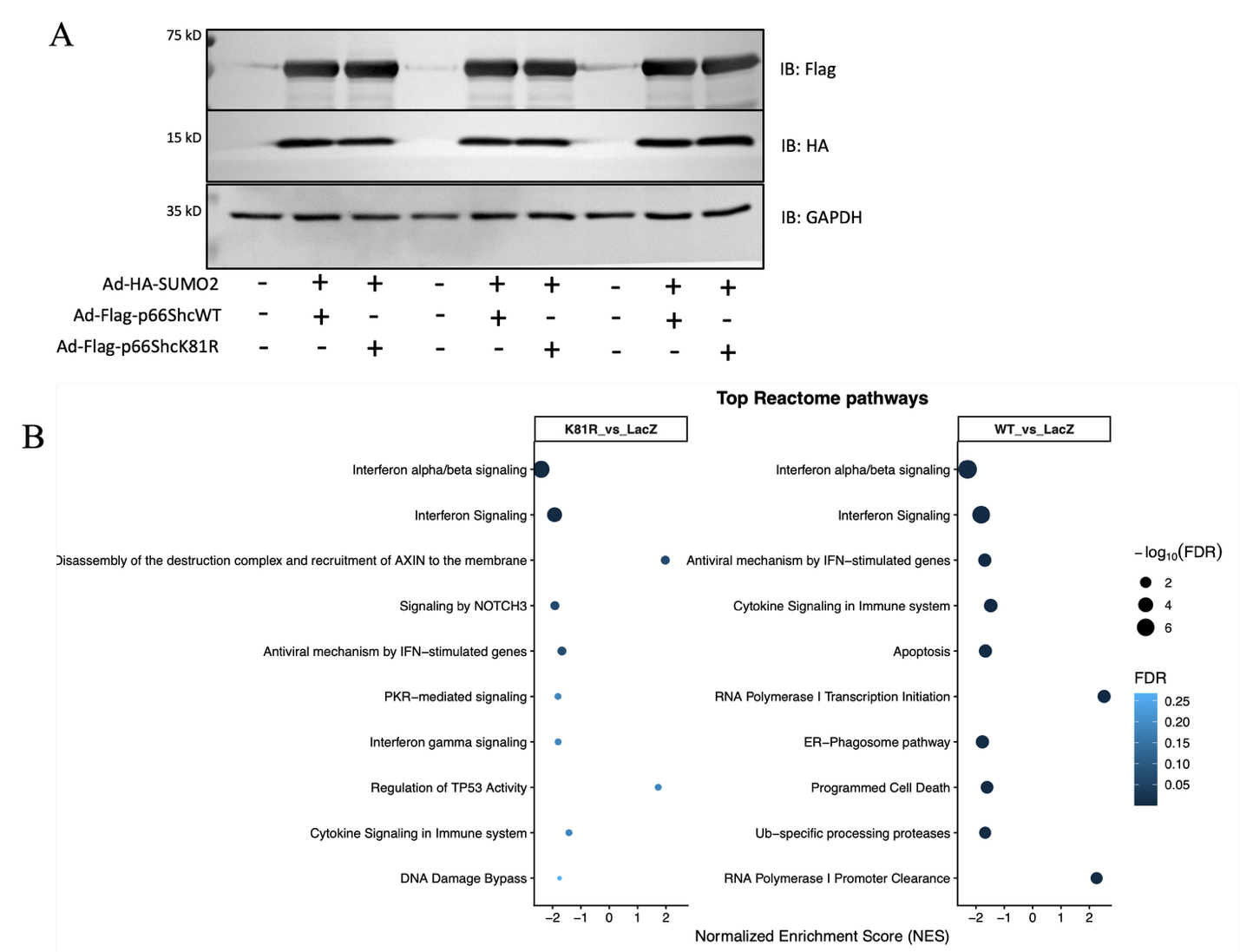

**Fig S5. P66Shck81-SUMO2ylation induced changes in signaling pathways in HUVECs.** (A) Representative immunoblots of endothelial cell lysate expressing Flag-p66ShcWT, Flag-p66ShcK81R, and HA-SUMO2 (n=3 shown, n=6 performed). (B) Bubble chart representation of all the regulatory pathways impacted by SUMO2-p66Shc in endothelial cells, rescued by p66ShcK81R. Pathway enrichment analysis (B) was performed using R-based Reactome pathway analysis and visualization packages on differential protein expression data obtained from HUVECs expressing SUMO2-p66ShcWT or SUMO2-p66ShcK81R compared with Ad-LacZ controls (n=6).

.
